# Joint estimation of paternity, sibships and pollen dispersal in a snapdragon hybrid zone

**DOI:** 10.1101/2024.01.05.574354

**Authors:** Thomas James Ellis, David Luke Field, Nicholas H. Barton

## Abstract

Inferring genealogical relationships of wild populations is useful because it gives direct estimates of mating patterns and variance in reproductive success. Inference can be improved by including information about parentage shared between siblings, or by modelling phenotypes or population data related to mating. However, we currently lack a framework to infer parent-offspring relationships, sibships, and population parameters in a single analysis. To address this we here extend a previous method Fractional Analysis of Paternity and Sibships to include population data for the case where one parent is known. We illustrate this with the example of pollen dispersal in a natural hybrid zone population of the snapdragon *Antirrhinum majus*. Pollen dispersal is leptokurtic, with half of mating events occurring within 30m, but with a long tail of mating events up to 859m. Using simulations we find that both sibship and population information substantially improve pedigree reconstruction, and that we can expect to resolve median dispersal distances with high accuracy.

## 2 Introduction

Knowledge of the genealogical relationships between individuals - their pedigree - is an extremely useful tool for understanding natural populations because it gives a direct estimate of who mates with whom, and differences in reproductive success (Pemberton, 2008; E. A. Thompson, 1976). One example, pedigrees can tell us about the distribution of dispersal distances organisms or gametes travel during their lifetimes, which sets the scale at which populations vary (Cain et al., 2000; Kremer et al., 2012). Of particular interest is both the average distance travelled and the shape of the dispersal distribution (Cain et al., 2000; Nathan et al., 2012). In plants, dispersal is often characterised by leptokurtic or ‘fat-tailed’ distributions, where dispersal is most likely to occur over short distances, but there is a long tail of long-range dispersal events (Austerlitz et al., 2004; Bullock et al., 2017; Clark, 1998). This leptokurtosis allows much more rapid dispersal than would be suggested by the average dispersal distance alone, with long-range migrants having a disproportionate effect on the spread of adaptive alleles (Cain et al., 2000; Clark, 1998). We thus aim to reconstruct mating events between individuals to infer the distribution of dispersal.

Pedigrees are inferred using three sources of information. First, the oldest and most common approach is to identify parent-offspring relationships that are most likely given on Mendelian inheritance, treating offspring individuals separately (Marshall et al., 1998; Meagher, 1986; E. A. Thompson, 1976). Second, sibling-reconstruction approaches use information about alleles shared between siblings to jointly infer sibling relationships with the parentage of the entire sibship (e.g. Anderson and Ng, 2016; Emery et al., 2001; Huisman, 2017; Jones et al., 2007; Thomas and Hill, 2002; E. Thompson and Meagher, 1987; Wang, 2004). Because sibling relationships are typically not known and there are a very large number of possible configurations to consider, a major challenge in sibship inference is to explore this space effectively. Genetic information shared between parents, offspring and between siblings are vital for reconstructing their relationships.

Finally, mating patterns and differences in mating success often depend on additional population and phenotypic parameters that are not related to genetics. Jointly modelling these processes with pedigree relationships can not only better resolve relationships between individuals, but provide more accurate estimates of population and phenotypic factors compared with post-hoc estimates (Hadfield et al., 2006; Neff et al., 2001). Pollen dispersal is a good example of this. We expect nearby individuals to mate with one another more often than with more distant individuals. By using a suitable model for how the probability of mating decays with distance we expect to weight inference to more likely candidate mates, but also to directly infer meaningful parameters about pollen flow (Adams et al., 1992; Chybicki & Burczyk, 2013; Jones, 2003; Klein et al., 2008; Oddou-Muratorio et al., 2005). A particularly well-established approach for this the the ‘neighbourhood model’ which jointly estimates parent-offspring relationships and dispersal distributions whilst also accounting for pollen immigration, incomplete sampling, variation in fertility and assortative mating (Adams & Birkes, 1989; Burczyk & Chybicki, 2004; Chybicki, 2018; Chybicki & Burczyk, 2013; Chybicki et al., 2021; Gérard et al., 2006; Oddou-Muratorio et al., 2005). Jointly modelling parent-offspring relationships with population parameters has great appeal because it allows us to directly address biologically relevant relationships between traits of interest and mating patterns.

Approaches exist for combining parentage with sibship information, and parentage with population parameters, and this additional information is expected to increase the accuracy of pedigree reconstruction (Neff et al., 2001; Walling et al., 2010; Wang, 2007). However we currently lack a framework for utilising all three sources of information parentage, sibships and population parameters - in a single joint analysis. Because of this, we also lack an understanding of the relative contribution of sibship and population parameters to pedigree inference, and how this changes depending on the data available and the biology of the system. We previously described a package Fractional Analysis of Paternity and Sibships (FAPS) to jointly infer paternity and sibships when the identity of one parent is known (Ellis et al., 2018). In this paper we describe how to include non-genetic information into the FAPS procedure to jointly infer paternity, sibships and population parameters. We then apply this to the inference of the pollen dispersal distribution in a hybrid-zone population of the snapdragon *Antirrhinum majus* with a large number of potential pollen donors. Using simulations we also characterise how the power to reconstruct paternity of individuals and identify mating pairs depends on the information available from genetics markers, the degree of shared paternity, and the scale of pollen dispersal.

## 3 Materials and Methods

### 3.1 *A. majus* data

#### 3.1.1 Study population

We examine a hybrid zone population of the snapdragon *Antirrhinum majus* ssp *majus* in the Spanish Pyrenees. Here, the yellow-flowered *A. m. majus* var *striatum* and the magenta-flowered *A. m. majus* var *pseudomajus* meet and hybridise to produce diverse recombinant phenotypes, including pink, white and orange flowers (Whibley, 2004). The population grows along two parallel roads running East-West close to Ribès de Freser (figure 1). The ‘lower’ road is at 1150-1200m above sea level, whilst the ‘upper’ road climbs 1250-1500m, and is 500-1000m north of the lower road. Hybrids are mostly confined to a 1km ‘core’ hybrid zone, with *A. m. striatum*-and *A. m. pseudomajus*- like plants becoming dominant to the West and East respectively. We surveyed as many flowering plants as we could find in June and July of 2012 (n=2124), and collected information on flower number and location using a Trimble GeoXT datalogger. We collected two to three leaves for DNA extraction and dried these in silica gel (Fisher Scientific). We also sampled one mature flower from each plant and scored this for yellow and magenta pigmentation as described by Whibley (2004). *A. majus* grows in disturbed habitats such as roadsides and railways; they are rare in the established forest and pasture between the two roads, on the north-facing slope to the South, and on the high mountain peak to the North of the two roads (figure 1). It thus is likely that we sampled the majority of the plants that flowered during the study period, although we cannot exclude that some flowering plants may have been missed.

**Figure 1.**
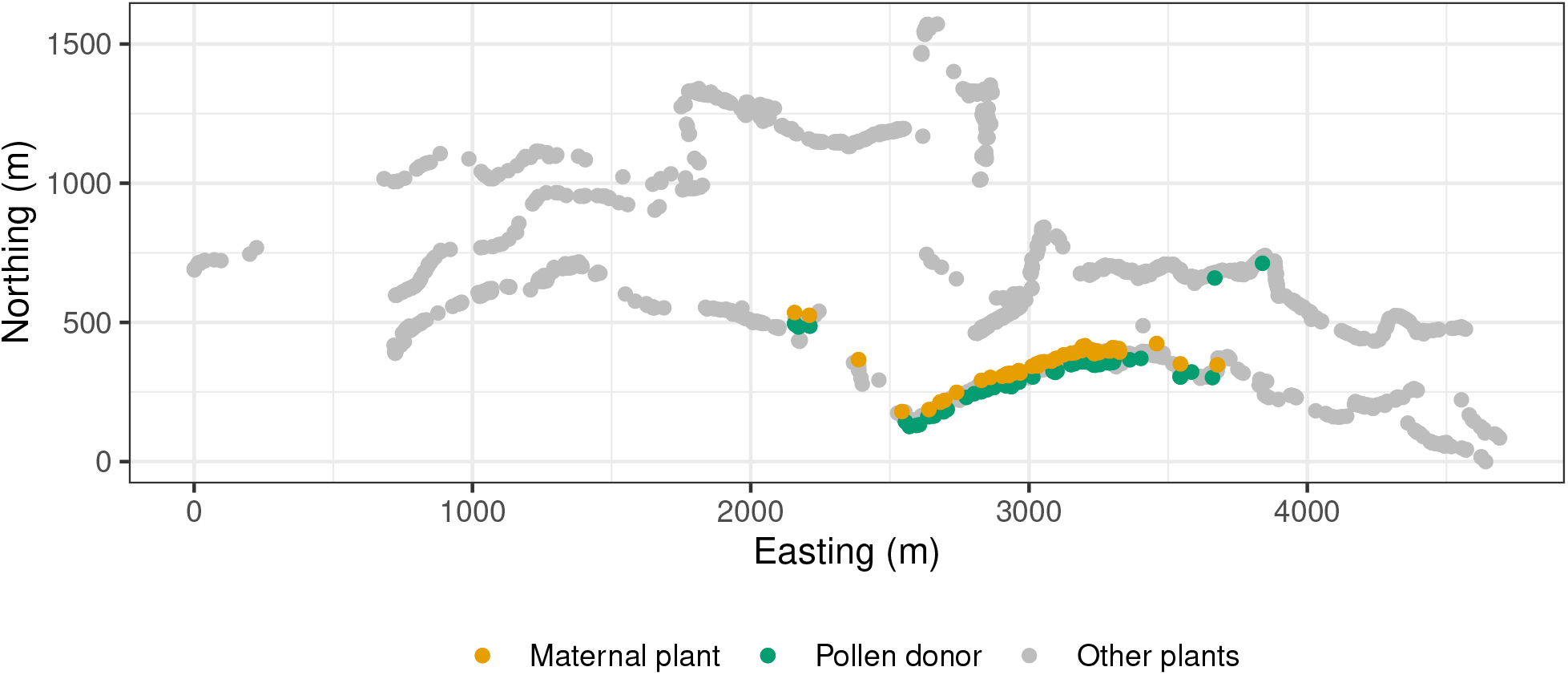
Map of the hybrid zone. The map shows the distribution of maternal plants, inferred pollen donors and remaining plants along the lower (South) and upper (North) roads. Only pollen donors for families with one or more offspring are shown.

Pollination is carried out exclusively by large bumble-bees and carpenter bees who are large enough to open the flowers (Andalo et al., 2019; Vargas et al., 2010). *A. majus* has a gametophytic self-incompatibility system, and self-pollinated seeds are very rare (Surendranadh et al., 2022). There are no detectable post-zygotic barriers (Andalo et al., 2010).

In August 2012 we collected a single, mature, wild-pollinated fruit from each of 60 mothers (figure 1). In order to minimise disturbance to the population we only sampled from plants which had set a minimum of five mature fruits. These mothers were chosen to represent an even sample of pigmentation phenotypes, spread as evenly as possible across the core of the hybrid zone where hybrids are most dense, resulting in 10 *A. m. striatum*-like, 17 *A. m. pseudomajus*-like, and 33 hybrid-phenotype mothers.

#### 3.1.2 Genotyping

We grew seeds in 5cm plug trays filled with potting compost (Gramaflor) in a greenhouse under Sylvania GroLux lights on a 16-hour cycle. We sowed three seeds per plug for 50-70 plugs per maternal family and thinned seedlings to a single seedling per plug after cotelydons had appeared. We transferred approximately 1cm^2^ of fresh tissue from 1419 seedlings to 96-well DNA-extraction plates (LGC Genomics, Berlin) and allowed tissue to dry using the sample bag and silica gel provided. For parental tissue from the hybrid zone we transferred approximately 1cm^2^ tissue dried in the field to the same plates. DNA extractions of the plated tissue samples were carried out by LGC Genomics.

We genotyped tissue samples at 71 SNPs by KASP sequencing (LGC Genomics). These SNPs are a sub-sample of a panel used for a wider survey of the hybrid zone (Surendranadh et al., 2022). Previous work identified the per-locus genotyping error rate in these data to be approximately 10^*−*4^ (Surendranadh et al., 2022). We removed 110 offspring that had missing data at more than 20% of the SNPs. We also pruned 4 SNPs that showed more than 20% missing data, or less than 15% heterozygosity. This left us with a set of 1308 offspring from 60 maternal families and 2124 candidate pollen donors, with between three and 39 offspring per maternal family (mean=21.8), with 64 marker SNPs.

### 3.2 Joint estimation of paternity, sibships and dispersal

#### 3.2.1 Probability model

We begin with observed SNP marker data **M** for mothers, offspring and candidate father, and a matrix **D** of Euclidean distances between mothers and all possible candidate fathers, and the per-locus genotyping error rate *ϵ*. From this we wish to infer pedigree *P* describing sibling, paternal and (known) maternal relationships, and vector *θ* of dispersal parameters. The full probability model is then

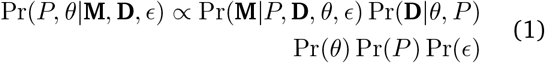

In the following sections we outline how to extend our existing method for inferring sibships and paternity to include data from non-genetic covariates, a model for pollen dispersal, suitable hyperprior distributions for dispersal parameters, and a procedure to infer the posterior distribution of the parameters of interest.

#### 3.2.2 Allowing for covariates in paternity inference

We have previously described a Python package FAPS which performs joint analysis of paternity and sibship relationships for sibling arrays based on SNP data, and allows for integrating out uncertainty in relationships (Ellis et al., 2018). Here we extended the software to allow for additional non-genetic information to be included.

FAPS begins with marker data for one or more maternal families composed of a mixture of half-and full-siblings, the known mother of each family, and an array of candidate fathers. Ellis et al. (2018) described a method to infer sibling and paternity relationships in those individuals; we refer the reader to that paper for the full details, but review the most important aspects here. FAPS uses marker data **M** to build matrix **G** of probabilities of paternity based on Mendelian transition probabilities for each maternal family. **G** has a row for each offspring and a column for each candidate father, where element *g*_*ij*_ is the probability that candidate *j* is the father of offspring *i* given marker data, and an estimate of genotyping error rate *ϵ*. The final column of **G** is the probability that the father of each offspring was missing from the sample of candidates, based on the probability of observing offspring alleles from population allele frequencies. Rows in **G** sum to one, and describe a multinomial distribution of probabilities of paternity over all candidate fathers, including unsampled fathers. FAPS then builds a similarity matrix whose *ih*^th^ element is the likelihood

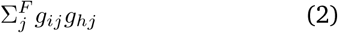

that the *i*^th^ and *h*^th^ offspring are full siblings by summing over the probabilities that they share any one of the *F* candidate fathers. The similarity matrix is used to perform hierarchical clustering to identify plausible ways to partition offspring into families of possible full sibships, and the likelihood of each partition structure is estimated by Monte-Carlo simulation. The likelihood that candidate father *j* is the father of putative full sibship *k* is then

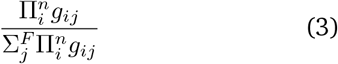

for all *n* offspring in *k*. FAPS returns a vector of probabilities (which sum to one) that each candidate is the father of sibship *k*, or that the true father was not sampled. This is done for each plausible partition structure. This gives a distribution of possible partition structures and their likelihoods, and a distribution of paternity probabilities for each putative family within each partition structure.

We can incorporate non-genetic information about paternity using a suitable function relating those data into probabilities of paternity (Hadfield et al., 2006). In principle this can be done for any kind of data for which there is a suitable function relating the observed data for a set of candidate fathers to a probability of having mating with each mother, given a set of parameter values for that function. The goal is to find parameter values that best explain the data. For example, a standardised continuous phenotype *z* could be modelled with the logistic function 1*/*(1 + *e*^*βz*^), where *β* describes the relationship between *z* and male fertility. A categorical phenotype can be modelled as a multinomial vector of probabilities that sum to one. In this study we use matrix **D** of Euclidean distances between mothers and candidate fathers as a covariate, and model the probability Pr(*d*^*mj*^|*θ*) of a mating event occurring between mother *m* and candidate *j* who are distance *d*_*mj*_ apart, based on a suitable model of pollen dispersal (see below). Assuming covariates of mating patterns are independent of marker genotypes it is then straightforward to incorporate this into the procedure outlined above by modifying eqn. 2 to

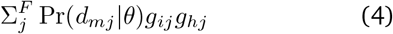

and eqn. 3 to

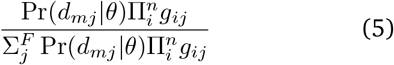

Eqn. 5 can then be used to estimate the likelihood of the partition structure by Monte-Carlo simulation, as previously described by Ellis et al. (2018). For a given set of dispersal parameters in *θ* the likelihood of the whole maternal family is then the sum of likelihoods for each possible partition. When there are multiple maternal families, the likelihood of the whole dataset given *θ* is the product of those likelihoods over each maternal family. When we update *θ*, this likelihood will change, giving us a way to compare likelihoods of different dispersal parameter values and identify those most consistent with the data.

#### 3.2.3 Pollen dispersal distribution

We wish to describe the distribution of distances travelled by pollen from anther to stigma (Nathan et al., 2012). A useful function for describing plant dispersal distributions is the generalised normal distribution (Clark, 1998; Kremer et al., 2012; Nadarajah, 2005). This is a generalisation of the exponential family of probability distributions and includes the exponential and standard normal distributions as special cases, but allows for fat and thin tails. It is commonly used to model plant dispersal distributions because these are often found to show clear kurtosis (e.g. Austerlitz et al., 2004; Burczyk et al., 2019; Field et al., 2011; Klein et al., 2008; Ottewell et al., 2012; Robledo-Arnuncio and Gil, 2005). The generalised normal distribution describes the probability of observing dispersal distance *d*_*mj*_ between mother *m* and candidate father *j* given scale parameter *a* and shape parameter *b* as:

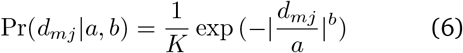

Where

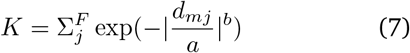

The function takes the forms of the standard exponential when *b* = 1 and normal distribution when *b* = 2. Values of *b <* 1 reflect leptokurtic distributions with tails that decay more slowly than would be expected under the exponential distribution. Using *θ* = {*a, b*}, this provides a convenient function for calculating Pr(*d*_*mj*_ |*θ*) because it allows for a long tail of long-distance migrants. Scaling by *K* ensures that probabilities of dispersal distances across fathers sum to one.

Sometimes it may be that an unrelated candidate has a similar or higher probability of paternity of one or more offspring simply due to stochasticity in Mendelian sampling. Whereas true fathers are expected to be (on average) close to the mother, other candidates should be drawn at random from the population (although this is complicated if there is substantial spatial genetic structure). This will inflate the apparent kurtosis in the data and bias *b* downwards. To accommodate this we modify eqn 6 to describe dispersal as a mixture of a generalised-normal and a uniform distribution:

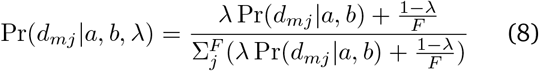

where *F* is the number of candidate fathers and *λ* is a mixture parameter determining the proportion of the probability mass due to ‘real’ dispersal. The uniform part of this mixture allows for signal coming from incorrect candidates without requiring the ‘true’ dispersal kernel to be unnecessarily leptokurtotic. This approach is conceptually similar to those of Streiff et al. (1999) and Slavov et al. (2009) who modelled dispersal as a mixture of distinct processes describing short-and long-range dispersal up to a maximum distance estimated from the data.

It is common to also report the mean dispersal distance as the square root of the variance of this distribution as

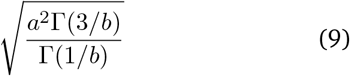

(Nadarajah, 2005). We prefer to focus on median dispersal distances because this is more intuitive for highly skewed distributions, and also to estimate this on realised inter-mate distances rather than on parameters alone (see “Inference of mating patterns”). We nevertheless report this and how it relates to the effect of *λ* on results.

#### 3.2.4 Priors for dispersal parameters

We require prior distributions for dispersal parameters *a, b* and *λ*. Because the effect of *a* depends on *b* and has no intuitive biological interpretation itself, we used prior simulations to choose a suitable parameterisation for *a* and *b*. We first simulated 10000 pairs of values for *a* and *b* from Gamma and log-normal distributions respectively. We then used each pair of simulated values to parameterise a generalised normal distribution, and simulated 1000 dispersal distances from that distribution, and visually examined the distribution for biological plausibility. We chose a gamma prior *a*∼ Γ(2, 100) for the scale and log-normal prior ln *b* ∼ *N* (*µ* = −0.7, *σ* = 0.5) for the shape parameters of the generalised-normal distribution. This model allows for a range of shape values reflecting leptokurtosis to Gaussian dispersal, and allows for long-distance dispersal events (figure S1). We also tried other prior distributions for *a* and *b* and found that they had very little influence on the posterior distributions (not shown).

For *λ*, we use a beta distribution with parameters Beta(1.1, 1.1). This distribution approaches zero when *λ* is close to zero or one, but is fairly flat in between. This implies that we do not expect that all the weight should be on either the generalised-normal or the uniform components of the mixture distribution in eqn. 8, but that we do not have strong prior beliefs about values between that. To examine the effect of modelling dispersal as a mixture model at all, we also repeated the analysis with *λ* to set to one.

### 3.3 Inference

#### 3.3.1 Inference via MCMC

We used the Metropolis-Hastings Markov-chain Monte Carlo (MCMC) algorithm to infer the posterior distribution of dispersal parameters *a, b* and *λ*. We ran four independent chains beginning from distant areas of the parameter space. At each iteration, we perturbed each parameter by a value drawn from a normal distribution with a fixed standard deviation for each parameter (*σ* = 2 for *a*; *σ* = 0.05 for *b*; *σ* = 0.025 for *λ*). We ran each chain for 25500 iterations and subsequently removed the first 500 iterations of each as burn-in. After checking chain convergence we thinned subsequent iterations to retain 250 posterior draws from each chain for further analyses, for a total of 1000 posterior draws.

#### 3.3.2 Inference of mating events

We aimed to create a list of possible mating events between mothers and candidate fathers consistent with the data to identify a set of independent pollen dispersal events. FAPS generates one or more partition structures that a half-sibling family could be split into full-sibling families. For each partition structure, FAPS identifies valid set of distinct fathers for each full sibship, and estimates a likelihood for each father-sibship configuration. Each set of fathers reflects a hypothetical set of mating events, and the probability that a single father mated with the maternal plant is the sum of probabilities for each partition structure in which he sires at least one offspring. This gives a list of mating events, with a posterior probability that each occurred, including an entry for offspring whose father was missing. The posterior estimate of offspring number is the number of offspring for each father in each father-sibship configuration weighted by the probability of the mating event. Note that the number and size of sibships are not necessarily integers because estimates are weighted averages over plausible sibship partition structures. We inferred mating events in this way using eqn. 3 based on dispersal parameter estimates from each of 1000 iterations of the MCMC output.

We previously noted that when FAPS errs, it tends to split one individual in a sibship into a singleton family, but essentially never incorrectly merges unrelated sibships (Ellis et al., 2018). We aimed to identify correct mating events based on family sizes. Previous investigations of FAPS showed that the main source of error was individuals from a larger sibship group being falsely assigned to a singleton sibship, although true half-siblings are almost never merged as full siblings (Ellis et al., 2018). This predicts that those singleton families will have posterior-mean family sizes of less than one because the nominal number of offspring is one, but the posterior probability for the family is less than one. This idea is not necessarily intuitive, so to illustrate this, imagine three offspring (A, B and C) with two candidate fathers (X and Y) and that there are two plausible paternal-sibships configurations:

1. X is the father of A, B and C, with probability 0.6,
2. X is the father of A and B while Y is the father of C, with probability 0.4.

The probability that X mated with the mother is 0.6+0.4=1, but the probability that Y mated with the mother is 0.4. Weighted mean offspring numbers are (3*0.6)+(2*0.4)=2.4 for X and 1*0.5=0.5 for Y. We can be more confident that the mother mated with X than with Y whilst being agnostic about the paternity of individuals. Based on this idea, we distinguish between mating events with posterior-mean family sizes greater or equal to one, and less than one.

Finally, we estimated the relationship between the number of offspring in a maternal array and the number of paternal families therein. We first averaged paternal family size for each mother-father pair over iterations of the MCMC, then fitted a Poisson generalised linear model (GLM) of paternal family size on maternal family size (McCullagh and Nelder, 1989).

### 3.4 Simulations

We used simulations to determine the statistical power of the dataset and and how this depends on the information used.

We simulated mating events between each true mother and one or more sires drawn from the pool of 2124 genotyped adult plants. Each maternal plant mated with the same number of pollen donors as inferred in the empirical analysis to generate one or more full-sib families per mother. For simplicity we drew sires for each family based on the probability of mating with a mother under an exponential pollen dispersal distribution (i.e. *b* = 1 and different values for *a*; see below). We ignored mating events with posterior probabilities less than 0.9. We simulated offspring genotypes based on Mendelian segregation and added genotyping errors to adults and offspring genotypes with the observed genotype error rate of 10^*−*4^ per locus per individual. In this way simulated data reflects the the observed genetic and spatial structure of the *A. majus* hybrid zone.

We parameterised simulations by varying four parameters that are likely to affect inference of relationships. We aimed to use values that reflect cases where parameters are less informative, moderately informative, and more informative. To vary marker information we downsampled the number of observed genotyped markers to 40, 53 and 67. To vary the information about paternity shared between full siblings we varied he number of offspring sired by each father (1, 3 and 5 offspring). To simulate missing fathers we randomly selected 10%, 30% and 50% of true sires and removed them from the set of candidate fathers. We simulated dispersal with mean distances of 3m, 30m and 300m. We repeated each parameter combination 100 times. We note that it is now routine to generate data with many more markers than used here, and in a previous study we did simulate additional SNPs (Ellis et al., 2018). However, denser markers will be in stronger linkage disequilibrium, meaning they are no longer independent and existing likelihood calculations are no longer valid. As such, additional markers do not necessarily translate into increased statistical power in a straightforward way. Using the observed markers also allowed us to capture the genetic structure of the population.

We used simulated datasets to infer mating events using the same procedure as for the empirical dataset. We compared results based on mating events inferred using three sources of information. First, we used most likely mother-father-offspring trios inferred using Mendelian transition likelihoods (‘paternity only’). Second, we inferred mating events using information about both paternity and sibling relationships (as described by Ellis et al., 2018). Third, we combined information from paternity, sibling relationships and the shape and scale of the dispersal distribution together (this paper). To test the hypothesis that we can distinguish between reliable and dubious mating events based on family size we quantified the probability that an inferred mating event was correct separately for mating events with posterior-mean family sizes of one or more, or less than one. To investigate different datasets and analyses estimated real dispersal we compared true realised median dispersal distances in each simulated dataset to median dispersal from inferred mating events.

## 4 Results

### 4.1 Family sizes vary in *A. majus*

In the empirical *A. majus* dataset we identified an average of 195 full-sibling families for which a father could be positively identified across posterior samples (96% credible intervals (CI): 189, 199). 15% of families had posterior probabilities and posterior-mean family sizes less than one, showed substantial uncertainty across MCMC iterations (figure S2) and seem to be drawn at random from across the population (figure S3), indicating that these mating events are likely to be dubious. In contrast, the remaining 165 families showed strong posterior support with little variation between MCMC iterations (figure S2), and were clustered closer to the maternal plants (figure S3). These likely reflect robust independent mating events that can be used to infer a dispersal kernel, and we focus on these mating events in the rest of this manuscript.

Full-sibship sizes ranged from one to 18 (figure S4). The GLM of the number of full sibships on maternal-family size revealed that (natural) log number of sibships increased by 0.047 for every additional offspring included in the maternal family (intercept = 0.092; figure S5). This corresponds to detecting a new family for approximately every 4 or 5 offspring genotyped, on average.

For all sixty mothers we found evidence for paternity by one or more unsampled fathers with non-zero probability (figure S2). These families include an average of 789 offspring (96% CIs: 785, 793). This corresponds to around 60% of the offspring.

### 4.2 Pollen dispersal is leptokurtic and asymmetric

The posterior distribution for the dispersal shape parameter was consistently less than one, with a posterior mean of 0.47 (96% CIs: 0.38, 0.58; figures 2A, S6), indicating a leptokurtic dispersal kernel. Posterior mean scale was 10.9 (96% CIs: 3.79, 22.4; figure 2B).

**Figure 2.**
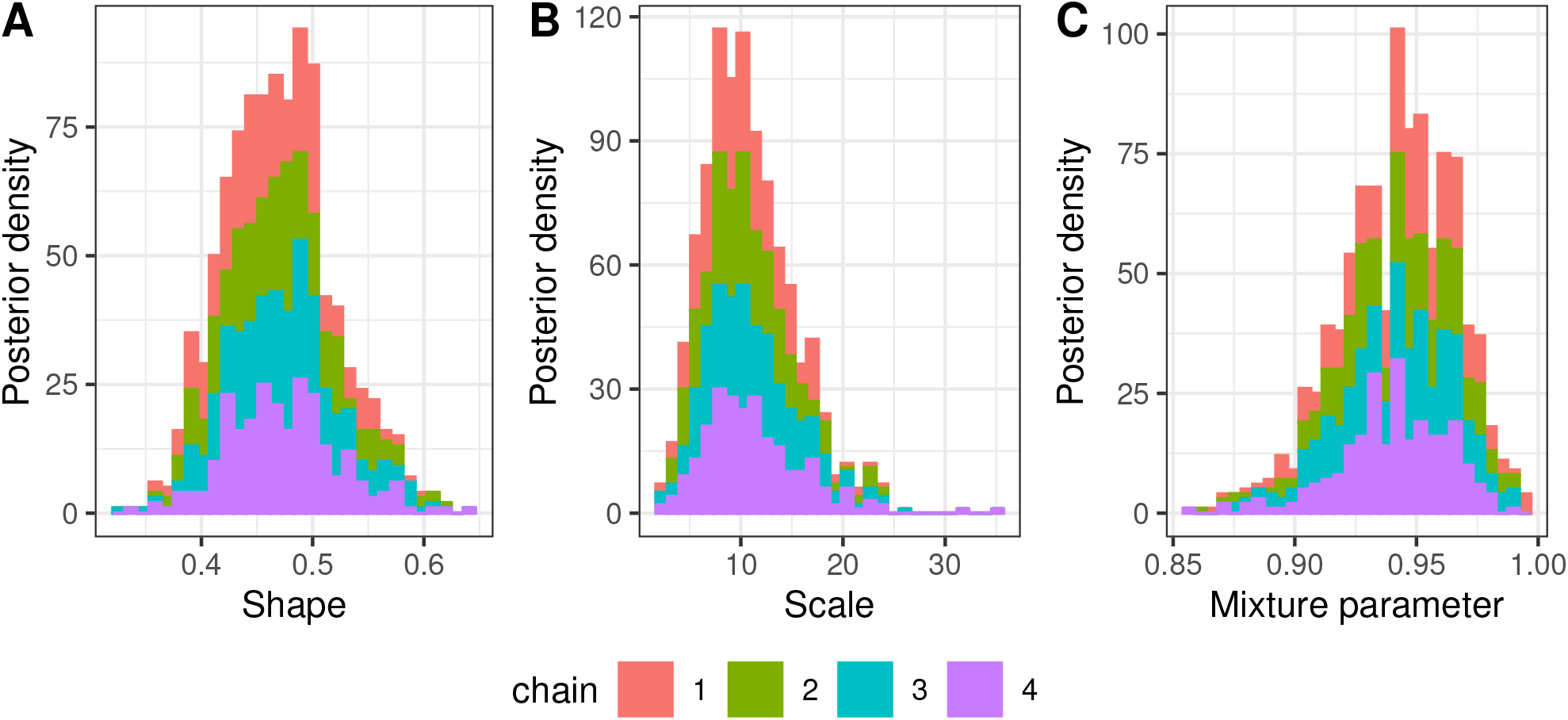
Posterior densities for the shape, scale, and mixture parameters of the dispersal kernel. Histograms show stacked densities for each of four independent chains.

Consistent with this, the distribution of pollen dispersal distances for families with one or more offspring shows a peak close to zero and a long tail of long-distance dispersal events up to 859m (figure 3). Mean and median dispersal distances were 83.4 and 28.0 metres respectively. With two exceptions, all mating occurred between plants on the lower road (figure 1). The cumulative distribution of realised dispersal distances varied very little across the posterior distributions of scale and shape parameters (figure 3B), indicating that the estimated dispersal distribution is robust. These results indicate a leptokurtic pollen dispersal kernel.

**Figure 3.**
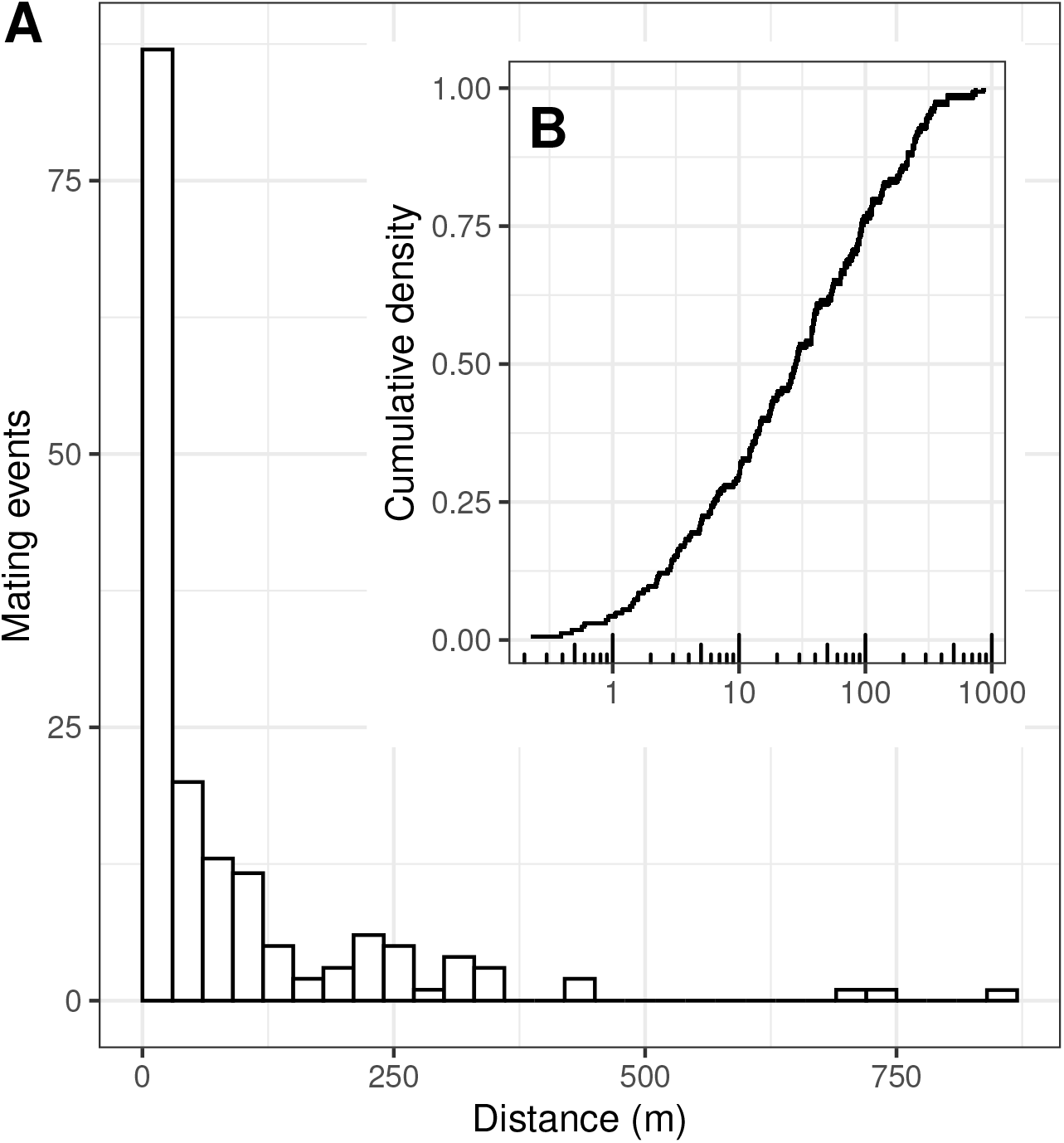
Distribution of pollen-dispersal distances. (A) Number of mating events in 30m bins. (B) Cumulative distribution of distances for dispersal events for paternal families with one or more offspring. Note the log_10_ scale of the x-axis in B. Mating events are weight weighted by their posterior probability. The histogram is summed over iterations of the MCMC. Separate cumulative curves are shown for 1000 MCMC iterations separately.

### 4.3 Mixture parameter influences dispersal shape

The posterior distribution the mixture parameter *λ* was centred around 0.94 (96% CIs: 0.89, 0.985; figure 2C). This indicates that up to 11% of paternal families are compatible with non-sires located at medium to long distances from the mother. This may be because the true father is missing and/or stochasticity in Mendelian sampling of the genotypes. If ignored, this signal would inflate apparent leptokurtosis in the dataset. Consistent with this, when we constrain *λ* to be fixed at 1 the posterior distributions for the scale and shape parameters are shifted downwards compared to the analysis where *λ* is allowed to vary (posterior means for *a* and *b* drop from 10.9 to 6.57 and from 0.47 to 0.39 respectively; figure S7). This would indicate more strongly leptokurtic pollen dispersal (figures 2A, 2B). When mean dispersal distance is estimated from the second moment of the generalised normal distribution (eqn. 9; Clark, 1998) this would imply a substantial increase in mean dispersal from 162 to 264 metres (figure S8). In this instance then, including a mixture component in the dispersal kernel reduces apparent mean dispersal distance when this is inferred from the moments of the generalised-normal dispersal distribution.

### 4.4 Sibship and population information improve pedigree and dispersal inference

We used simulations to investigate how well we can distinguish true and false mating events based on posterior-mean family size. When families contained only a single offspring a large proportion of correct mating events had dubious support and posterior-mean family size less than one (figure 4A). This was more pronounced with increased numbers of markers, and when dispersal information was included in the analysis. When real family sizes were greater than one however, almost no true mating events had posterior-mean family sizes less than one. This means that when families contain multiple siblings there is no cost to ignoring apparent singleton families. Distinguishing true and false mating events based on family size gives mixed results for singleton families, but is an effective means to identify true mating events for larger families.

**Figure 4.**
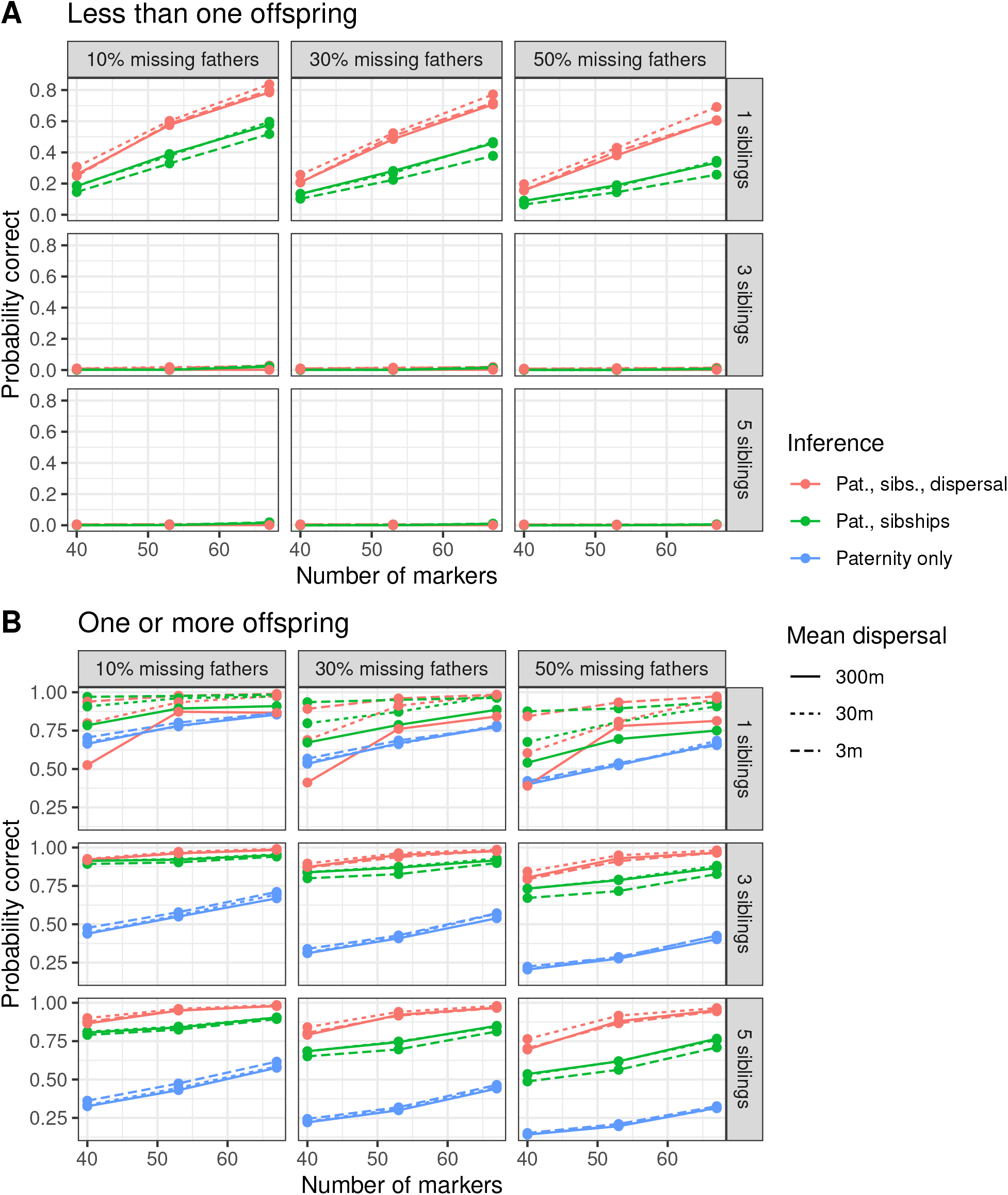
Mating events in simulated data. Values show the probability that inferred mating events are correct for families with posterior-mean family sizes of (A) more or more or (B) less than one. Panels show the proportion of true fathers missing from the dataset used to infer mating events (columns) and the true full sibship size (rows). Values are the mean of 100 replicate simulations, and are shown for inference using paternity (Pat) only, paternity and sibships (sibs), and paternity, sibships and dispersal information. Note that values for inference with paternity only are not shown in A because these genealogies are inferred from the most-likely configuration, so a posterior-mean family size is not defined.

The simulations also reveal how well we can reconstruct mating events under different scenarios and using different data. Inferred mating events were more likely to be correct as the number of marker loci and the proportion of true fathers that were sampled increased, and decreased with larger families (figure 4B). We found small differences in accuracy between dispersal scenarios, but the direction depended on what information was used to infer mating events. For example, when only paternity trios were used, the least informative scenario (mean dispersal = 300m) showed the highest accuracy, and the most informative (mean dispersal = 3m) the lowest, but when information from paternity and sibships were combined the opposite pattern was true. The clearest result is that there was a substantial gain in power to detect mating events when paternity and sibship information are used relative to paternity only, and when paternity, sibship and dispersal information are used relative to paternity and sibships only. Including information from both sibships and population parameters greatly increase the accuracy of inferred mating events.

A substantial proportion of simulated offspring were assigned to unsampled fathers (figure S9). Interestingly, this increased with increasing number of markers, indicating that FAPS becomes more conservative about assigning paternity to sampled fathers as marker information increases. We found that models using only paternity, or paternity and sibship information tended to underestimate the true proportion of offspring whose father was missing when that proportion was high, but were largely correct when that proportion was low. In contrast, the model that included dispersal information tended to overestimate the true proportion of offspring whose father was missing when that proportion was low, but were largely correct when that proportion was high. The improvement in identifying true mating events when including additional information may come with a cost of increased false-negative assignment to unsampled fathers in some circumstances.

We compared median dispersal of inferred mating events to true median dispersal (figure 5). Dispersal estimates were substantially more accurate when more markers were used, but showed less of a dependence on family size or the proportion of sampled fathers. Instead, estimated dispersal distances depended strongly on true dispersal distance; estimates were most accurate for datasets with the shortest dispersal and least accurate for dispersal over the largest distance. Estimated dispersal also depended strongly on the information used to infer mating events. Analyses using paternity information only often overestimated dispersal by more than a hundred metres. Including information about sibling relationships reduced this estimate substantially, and including information about dispersal reduced these even further. Of particular note is that when all 67 markers are used, as in our empirical dataset, the estimate of median dispersal is typically within one metre of the true value, indicating that our empirical estimate is robust. Inference of dispersal distance is most reliable when dispersal is over relatively short scales, and as much information as possible is included in the analysis.

**Figure 5.**
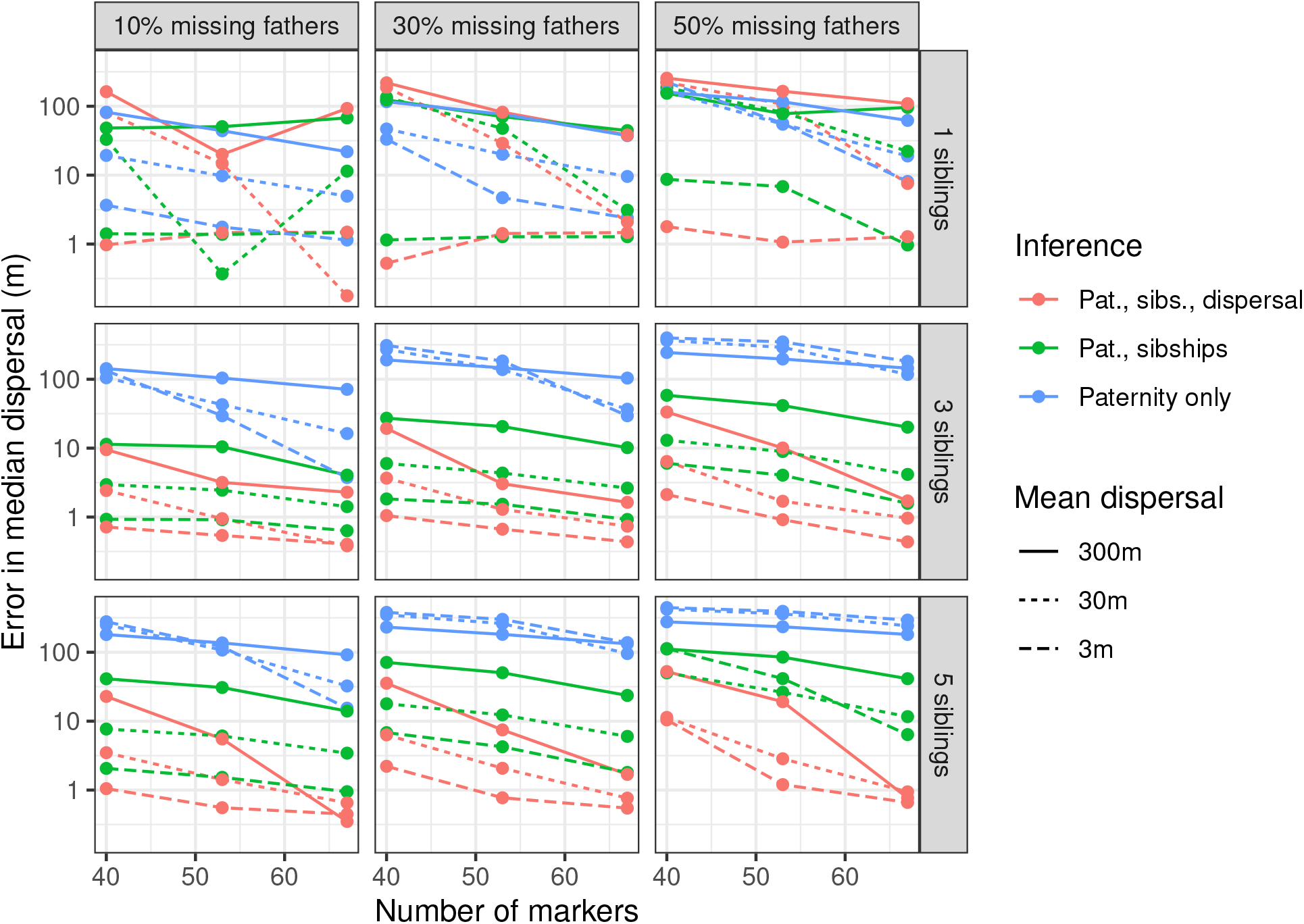
Errors in dispersal estimates in simulated data. Values show the absolute difference in median dispersal events between real mating events and inferred mating events with posterior-mean family sizes of one or more. Panels show the proportion of true fathers missing from the dataset used to infer mating events (columns) and the true full sibship size (rows). Values are the mean of 100 replicate simulations, and are shown for inference using paternity (Pat) only, paternity and sibships (sibs), and paternity, sibships and dispersal information.

## 5 Discussion

We have described a framework to jointly infer paternity, sibling relationships and population parameters. Applying this to a natural populations of snapdragons, we find strong evidence for a leptokurtic pollen-dispersal distribution. Using simulations we demonstrate that sibship and population information improves inference of genealogical relationships and biological conclusions.

### 5.1 Implications for the hybrid-zone population

The shape of the dispersal kernel implies that most mating occurs within tens of metres, but that there is a long tail of mating events between individuals at up to several hundred metres. This result suggests that any natural selection acting through male reproductive components (pollen dispersal) occurs at small spatial scales. If pollinators discriminate between different flower-colour phenotypes in the hybrid zone, this sets an important geographic scale of selection for empirical tests on pollinator-mediated selection. For example, within 20m in the core of the hybrid zone, pollinators would encounter a full range of colour phenotypes that would provide scope for colour discrimination and differential visitation to influence male and female fitness. In this paper we have not attempted to estimate selection via flower colour because the number of mothers for each flower colour is small, meaning that we likely have little statistical power. However, in ongoing work we are investigating variation in fitness in a much larger panel of offspring arrays, which will allow us to address such questions with much greater accuracy.

Rare long-distance dispersal events can have unexpectedly large effects on natural populations (Cain et al., 2000; Clark, 1998). We found that the most extreme pollen-dispersal events spanned many hundreds of metres. Since the width of the core of the hybrid zone is approximately one kilometre, this means that pollen would be able to quickly move across a substantial fraction of hybrid zone, and accelerate introgression. However, there are several reasons why long-distance dispersal events are especially difficult to detect. First, these events are intrinsically rare. Second, as distance increases the more plants need to be sampled to ensure the father is included. Third, as more candidate fathers are included the greater the chance that an unrelated fathers is falsely assigned. It is thus likely that we have underestimated the tail of long-distance dispersal events to some extent. However, because this population is mostly confined to roadsides, our population sample extends several kilometres further than the longest dispersal event detected, and we can minimise false-positive paternity assignments by conditioning on family size, this underestimate is unlikely to be substantial. Nevertheless, it remains possible that dispersal events over even greater ranges may occur in this population, for example by larger queen bumblebees early in the season (Lepais et al., 2010), with the potential to have a large impact on gene flow.

### 5.2 Controlling bias in dispersal estimates

#### 5.2.1 Bias due to incorrect fathers

Sometimes candidate pollen donors can have a similar or greater Mendelian probability of paternity for an offspring individual as does the true father due to stochasticity in Mendelian sampling (E. Thompson, 1976). If not addressed, we might infer an incorrect mating event between adult plants, which would bias estimates of dispersal upwards. Two aspects of our study suggest that signal from false fathers should have a minimal effect on our results.

First, our simulations showed that incorrect families could be readily identified based on family size. In previous simulations to test the performance of FAPS we found that the main source of error was individuals from a larger sibship group being falsely assigned to a singleton sibship, but this analysis focussed on the most likely partition structure to simplify results (Ellis et al., 2018). In this study we found that those incorrect singleton families have low posterior probabilities when considered as part of a whole set of plausible sibship configurations, meaning that family sizes are effectively less than one individual. Since our goal was to infer pollen dispersal by identifying mate pairs, excluding mating events for families with sizes less than one is an effective way to identify a conservative set of high-confidence mating events, even if this means that some true mating events are lost. Consistent with this, in simulated data that closely match the *A. majus* dataset we found that estimated dispersal was very close to the true values, indicating that false-positive mating events have not had a large influence on our estimates of dispersal. This demonstrates that accounting for uncertainty in sibship structure is an extremely useful way to improve the accuracy of paternity inference.

Second, we modelled dispersal as a mixture distribution with a term allowing for false-positive fathers to reduce the bias in estimates of dispersal. The mixture distribution reduces the bias in the estimated shape parameter and the second moment of the generalised normal distribution, often interpreted as mean (squared) dispersal distance (e.g. Austerlitz et al., 2004; Clark, 1998; Klein et al., 2008). We note that this bias will exist to some extent as long as knowledge of paternity is less than perfect. Moreover, two factors may complicate the interpretation of the mixture paramter. First, the idea assumes that unrelated candidates are distributed at random with respect to genotypes, which will not be the case if there is spatial genetic structure. In the case of the *A. majus* populaton the marker panel was designed to show very little spatial structure, so this is unlikely to have a substantial impact. Second, more distant mating events may be real, but reflect some other distribution process than that modelled by the generalised normal distribution. In this case, the mixture component is still useful in that it allows these two distributions to be decoupled, at least to some extent (Slavov et al., 2009; Streiff et al., 1999). Here, our biological conclusions are not affected, because we focus on the distribution of realised mating events, which are affected in only a minor way in real terms (figure 3). However, caution is warranted in the interpretation of the raw parameter values for shape and mean dispersal from the generalised normal distribution (figure S8). It is possible that the effect was stronger here than would be the case in other published studies because our sample of candidate fathers is large, and the spatial extend is much larger (two orders of magnitude) than median dispersal distance. In contrast, other studies have focussed on wind-pollinated trees, where pollen can travel much further, and had smaller numbers of candidates (e.g. Adams et al., 1992; Austerlitz et al., 2004; Klein et al., 2008). This demonstrates the utility of using inferred dispersal parameters to inform inference of real mating events, rather than focussing on the parameters themselves.

#### 5.2.2 Missing fathers

An alternative source of bias comes from false negative paternity, where the true father of a family is present but is inferred to be missing. We found that more than half the offspring were inferred to have an unsampled father. Since these are cases were paternity has not been falsely assigned to an unrelated candidate, we do not expect false assignment to bias dispersal estimates upwards. Nevertheless, the extent to which these missing fathers bias dispersal estimates still depends on their distribution in space. At the same time, it may be that sampling effort was greater in the core of the hybrid zone than at the edges of our sampling region. It is known that bumblebees, especially queens, can disperse over several kilometres, so very long range pollen dispersal is plausible (Hagen et al., 2011; Lepais et al., 2010; Osborne et al., 2008). In this way it is possible that we have missed long-distance mating events, which would bias dispersal estimates downward. However, our sample of candidate fathers is large, and extends at least a kilometer beyond the range of plants identified as pollen donors (figure 1), which is more than twice as far as the longest dispersal event detected (figure 3). We feel it is thus unlikely that there could have been enough unsampled fathers far from the sample of mothers to greatly influence dispersal shape or median dispersal distance.

An alternative explanation for the high number of missing fathers is that they finished flowering before the main sampling began. This is a more plausible explanation, because the population had already begun to flower at the time of the first surveys in June 2012, and plants were still flowering at the time of seed collection in late July. The majority of unsampled fathers may thus represent relatively early-flowering individuals. Fortunately, it seems that flowering time is unlikely to depend strongly on location, and as such this is unlikely to bias dispersal estimates.

### 5.3 The value of sibship and population data

We found that information about sibling relationships and information about dispersal improved identification of mating events in simulated data. In a similar study, Walling et al. (2010) compared parentage assignment in a wild deer population inferred from the programs Cervus (which uses parent-offspring trios; Marshall et al., 1998), Colony (which infers parentage and sibling relationships; Wang and Santure, 2009) and MasterBayes (which infers parentage and population parameters; Hadfield et al., 2006). Both Colony and MasterBayes showed a similar increase in paternity assignment when sibship and population data were included over analyses using the same programs that only used parentage information, but were not able to combine the sources of information. Here we include both sibship and population information into a single analysis and find that they are useful independent of one another.

There was surprisingly little difference in the accuracy of mating events with increasing family sizes in simulated data (figure 4B). This is counter to expectations by Wang (2007) that power should increase with more siblings. This may be partly due to the structure of the simulations. We kept the number of mating events fixed, so increasing family size means increasing the total number of progeny. That in turn means that there are more opportunities for an offspring to be incorrectly assigned to an unrelated father. In addition, FAPS estimates likelihoods of full sibships by multiplying probabilities of paternity for individuals, which under-utilises the available information about parentage shared between siblings (E. Thompson & Meagher, 1987). As such, it is possible that we have underestimated the value of sibling relationships in this study.

It is helpful to view population data as acting like a Bayesian prior that places different weights on paternity probabilities independent of genotype data (eqn 1). In the case of dispersal, an expectation that dispersal occurs over shorter distances favours candidates close to the mother, which effectively narrows the pool of plausible candidates, and limits the number of unrelated candidates who could incorrectly assigned as a father. This explains the observation that including dispersal information decreased the number of incorrect mating events and increased the proportion of offspring assigned to missing fathers (figures 4B, S9). It also explains why estimated median dispersal distances were more accurate for narrower dispersal distances (figure 5). Our simulations did not investigate the effect of how badly one can misspecify the shape of the dispersal kernel before results are untenable, which would be a useful future avenue.

In this paper we have outlined a general method to combine data about population parameters with analyses of paternity and sibship. We have modelled dispersal distances this as one such factor affecting mating probabilities to illustrate this, but this clearly underestimates the complexity of interesting mating patterns that could be modelled. For example, we would expect that a pollen donor’s overall floral display size, which may be viewed as a proxy for fecundity, as well as the extent to which flowering phenologies overlap to be positively correlated with the probability of mating with a maternal plant. Likewise, flower colour may well also play a role in mating, although it is less clear exactly how. Although it would be feasible to include such data to inform pedigree analyses (see Methods), we have not attempted this here because we currently lack the data to do this in a meaningful way. Ongoing work currently aims to collect much more detailed demographic and phenotypic information from many more plants, which will allow us to explore these ideas further.

## Supporting information

Text file detailing observed mating events

## 6 Acknowledgements

We thank a large number of field volunteers for maintaining the population sampling, and Tom White for assistance with seed collection. We thank Sylvia Rebel for plating tissue for DNA extraction, as well as Sean Stankowski and two anonymous reviewers for feedback on the manuscript.

## 7 Data availability

Data and code to recreate the analyses are available from Zenodo (Ellis, 2025). A supplemental file listing inferred mating events is included as a supplementary file.

### 8 Author contributions

TJE sampled seeds and processed seeds, designed the method, analysed the data and wrote the manuscript. DLF led the collection of the demographic data of the hybrid zone, designed the SNP panel. DLF and NHB provided critical feedback on the manuscript.

## Notes

https://doi.org/10.5281/zenodo.15657146

### Competing Interest Statement

The authors have declared no competing interest.

### Summary of Updates

We have modified the manuscript to expand the parameter range of simulations we consider, and assess the effect on accuracy of identifying correct fathers, correct mating events, and median dispersal distance. We have also modified the priors on the main MCMC analysis to be less skeptical about long-range dispersal. In doing so we modify the prior on the shape of the dispersal kernel to weight towards slightly more leptokurtic distributions.

