## Supplementary figures for "Joint estimation of paternity, sibships and pollen dispersal in a snapdragon hybrid zone"

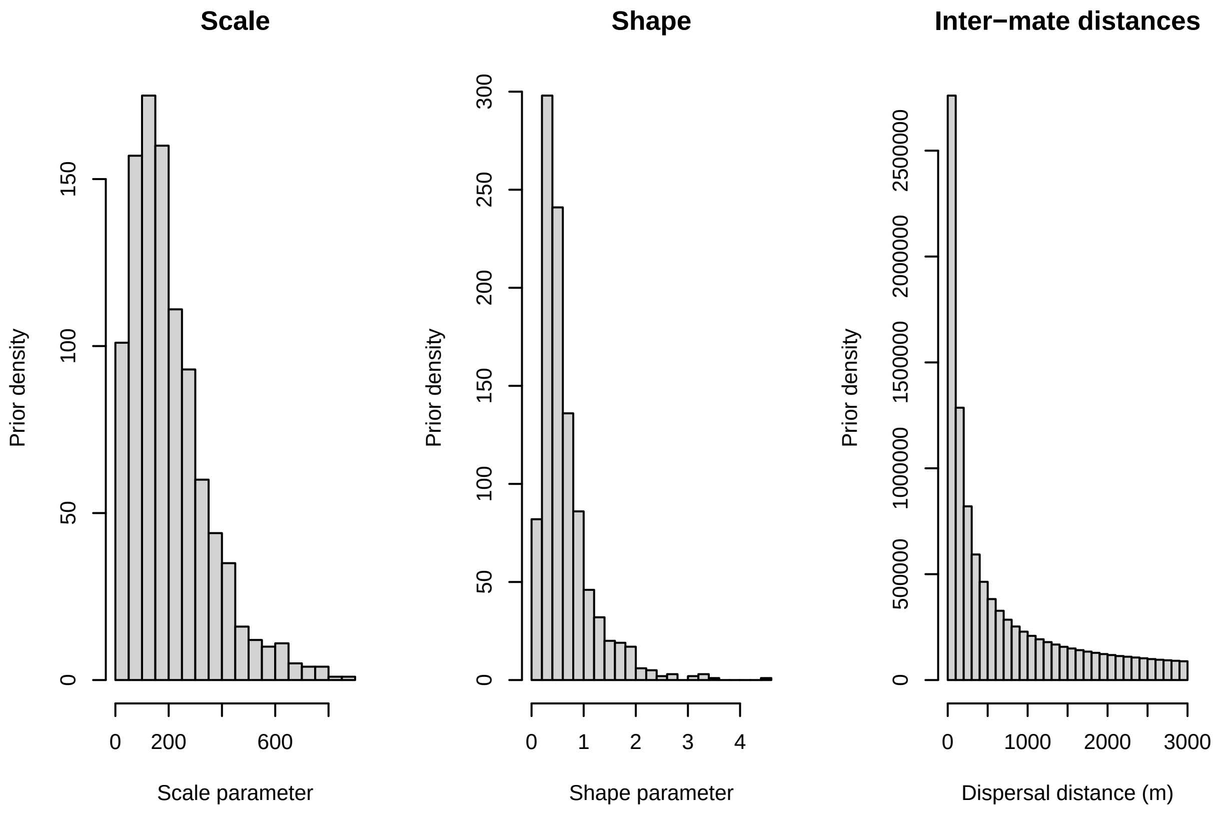


Figure S1: Prior simulations. Histrograms show simulated values for the scale (A-C) and shape (D-F) parameters of the generalised normal distribution, with and the dispersal distances those imply (G-I). Simulations are shown for three prior contrasting dispersal scenarios: (A,D,G) a model with moderate skepticism about leptokurtotic dispersal; (B,E, H) a model allowing leptokurtosis, but skeptical of long-range dispersal; (C,F, I) a model highly skeptical of leptokurtosis. In A-F, Gamma priors on the shape and scale of the GND are giving, showing the shape and rate of the Gamma distribution. D-F also indicate the proportion of the prior mass on leptokurtotic distributions (when $b<1$). Each histogram is summed over 1000 replicate draws from Gamma priors.

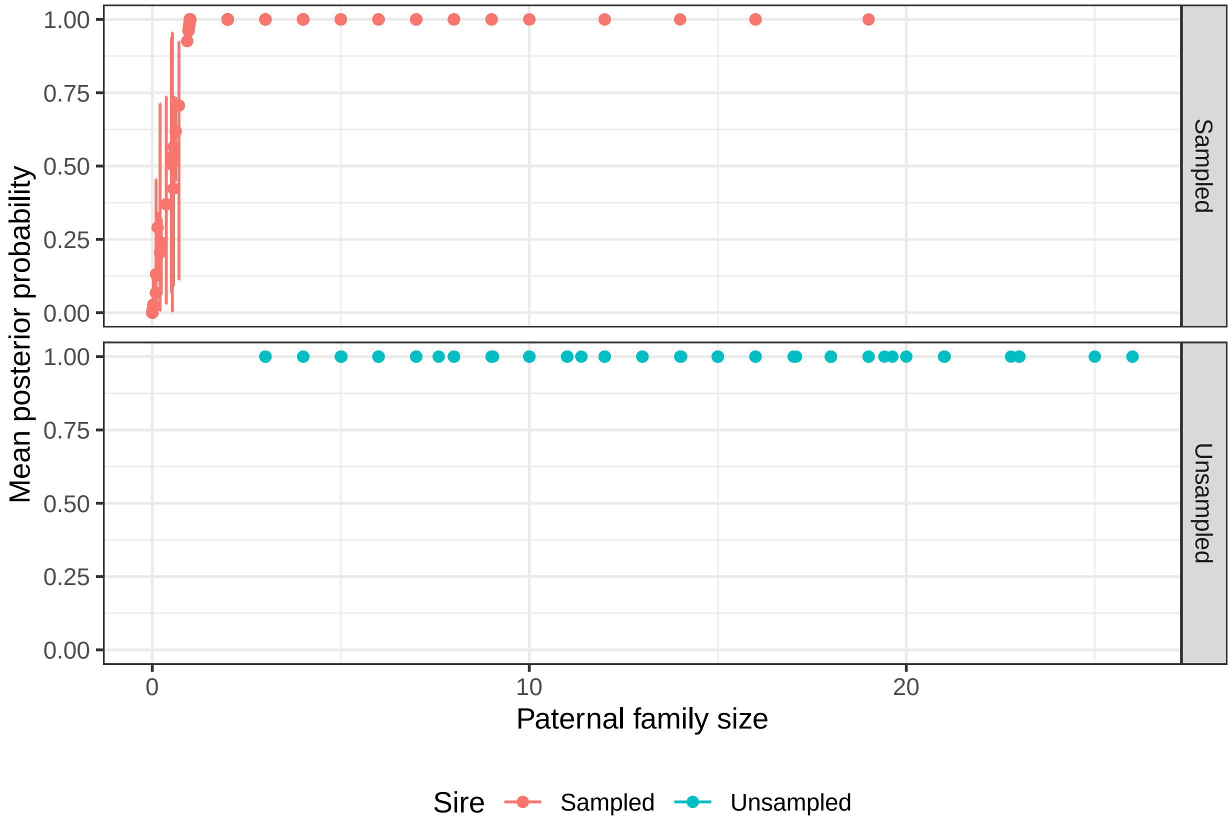


Figure S2: Posterior probabilities and sizes of paternal families. Each point shows the mean probability of a mating event between a mother and a single father averaged over 1000 MCMC iterations for the 'restricted-kurtosis' prior scenario, with 96% credible intervals. Colour indicates whether the father was in the sample of candidate or was unsampled. Note that individuals in families with unsampled fathers may consist of multiple paternal families.

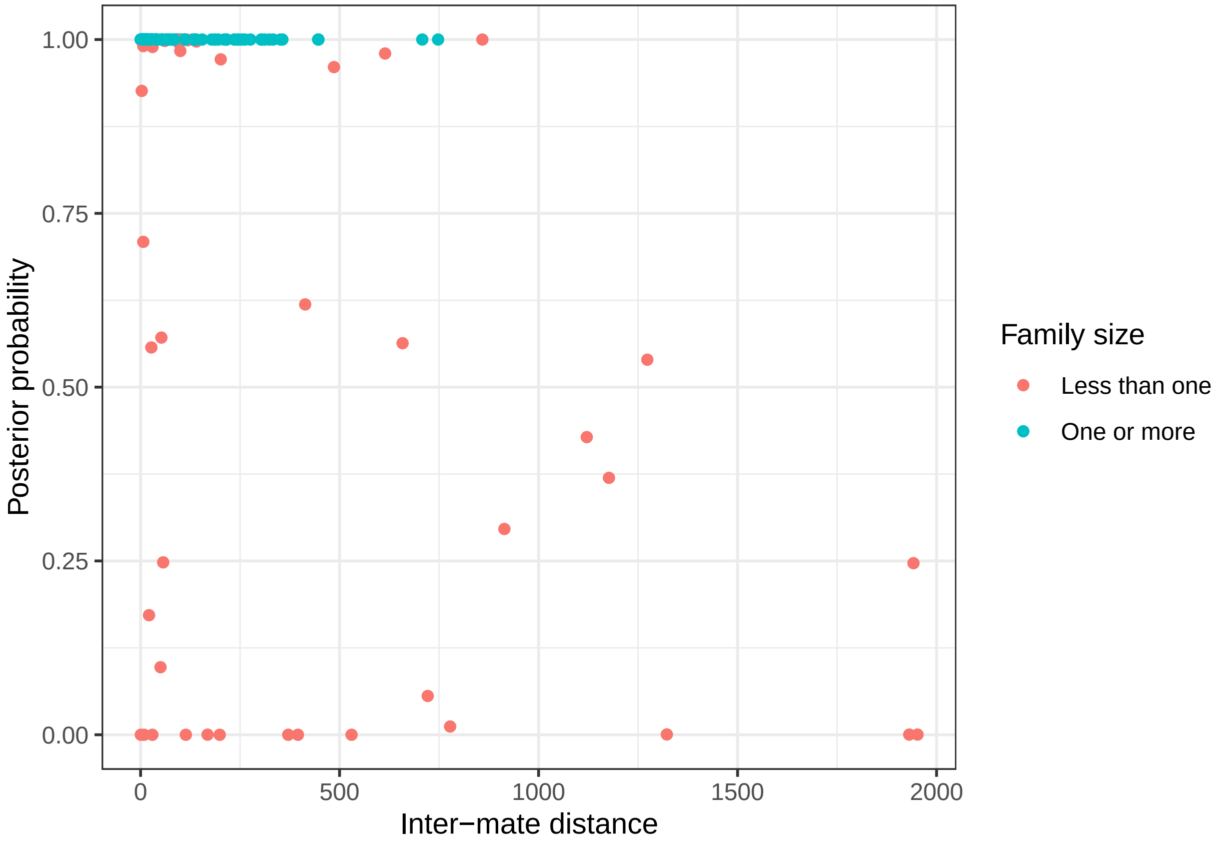


Figure S3: Relationship between the distance between mothers and inferred fathers and the posterior probability of the mating event. Points are coloured separately to indicate whether family sizes are less or greater than one.

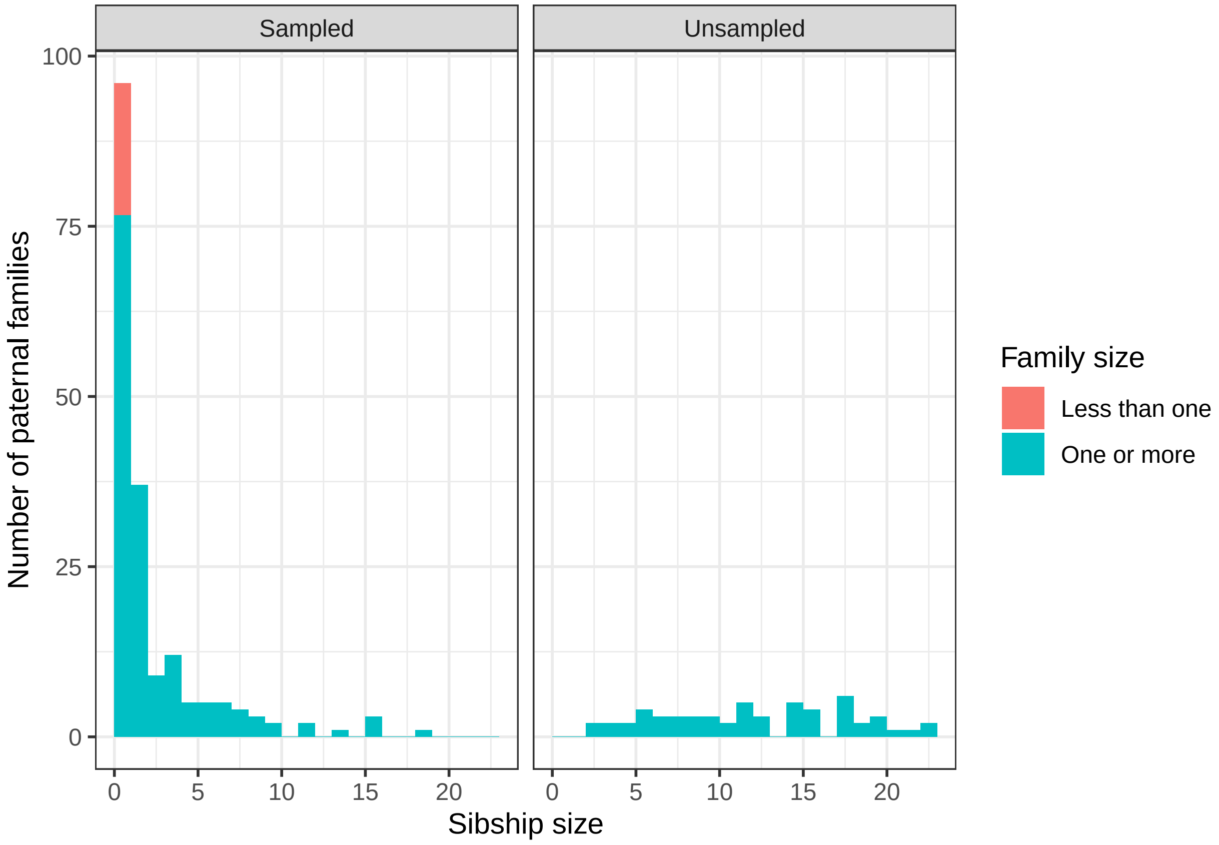


Figure S4: Histogram of paternal family sizes weighted by posterior probability of each family. Data are shown for families with an identifiable father, and those inferred to be sired by an unsampled father. Colours indicate whether weighted-mean family size was more or less than one.

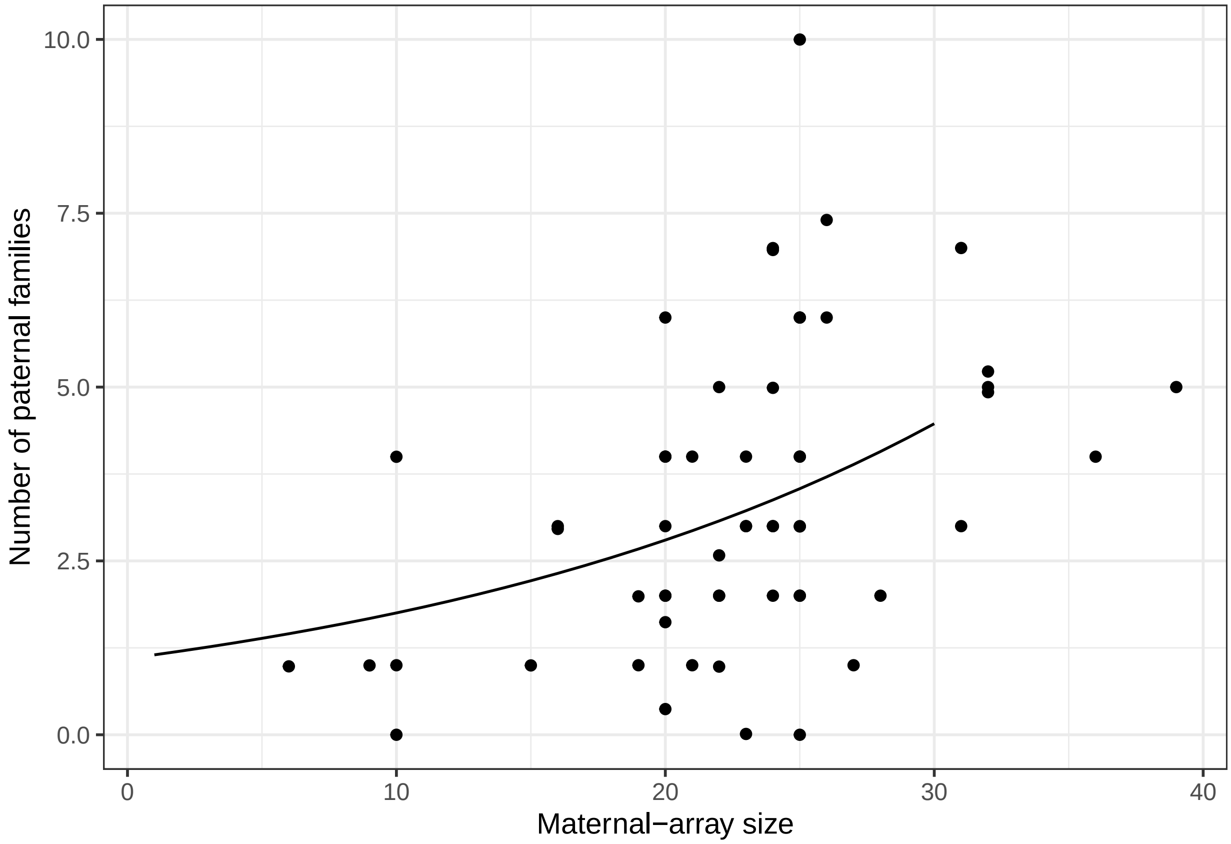


Figure S5: Number of paternal families as a function of total maternal array size. Data shown for the 'restricted-kurtosis' prior scenario and are averaged over 1000 iterations of the MCMC. Also plotted is curve predicted from a Poisson GLM of family number on array size, constrained to go through zero at the intercept.

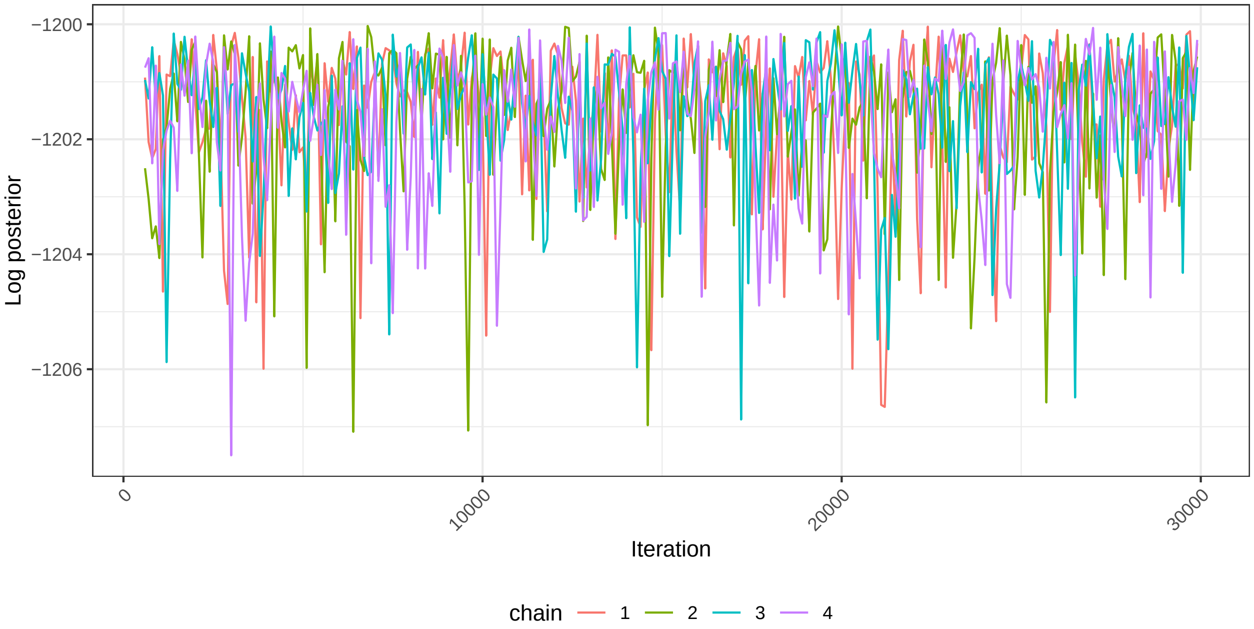


Figure S6: Change in log posterior density over four independent MCMC chains. The first 500 iterations of each chain have been removed as burn-in.

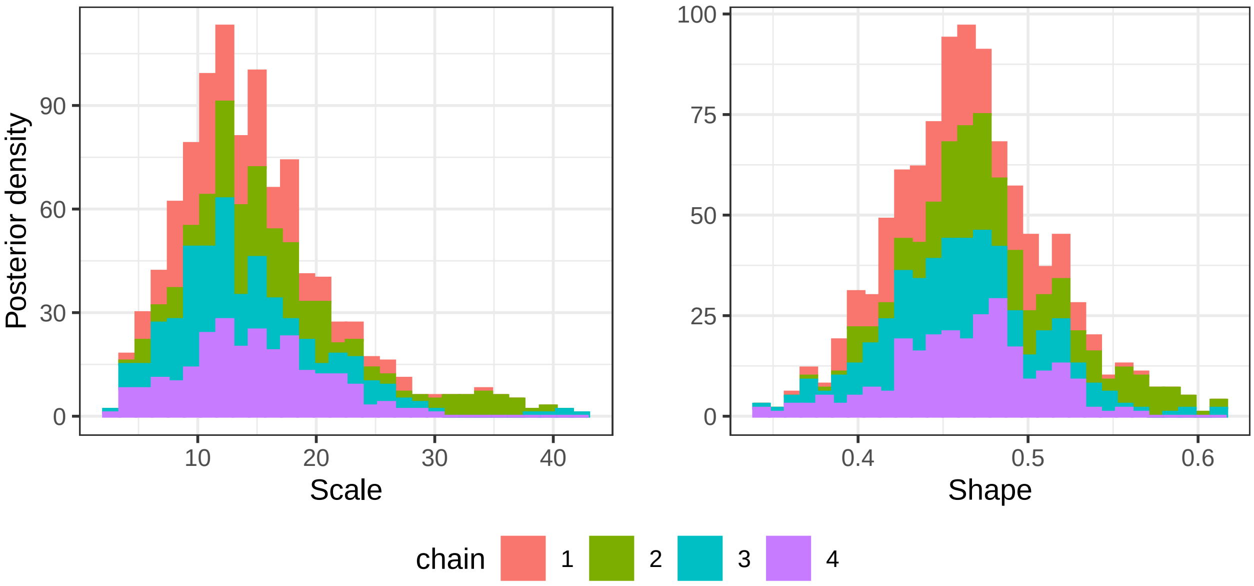


Figure S7: Posterior densities for the shape and scale parameters of the dispersal kernel for a model where the mixture parameter is fixed to one. Histograms show stacked densities for each of four independent chains.

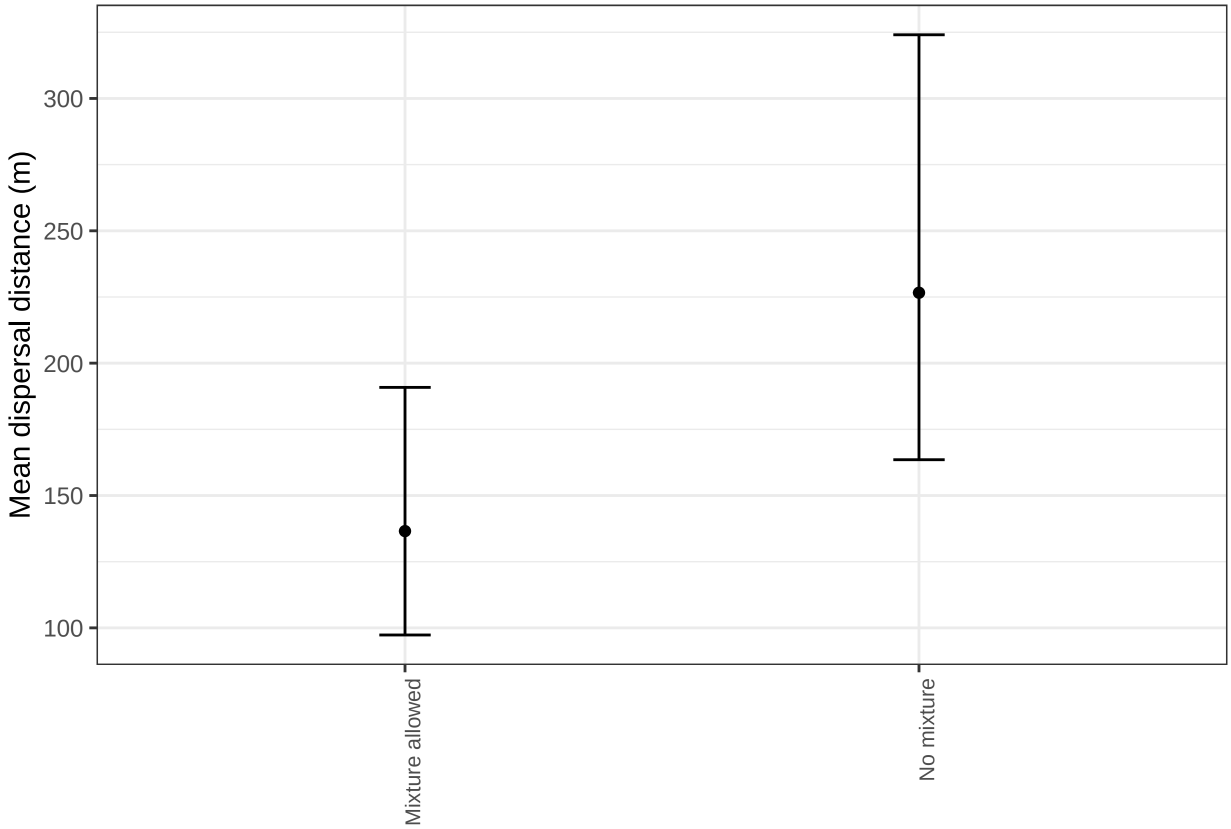


Figure S8: Mean dispersal distances implied by the second moment of the generalised normal distribution based on scale and shape parameters from MCMC chains for six prior scenarios.

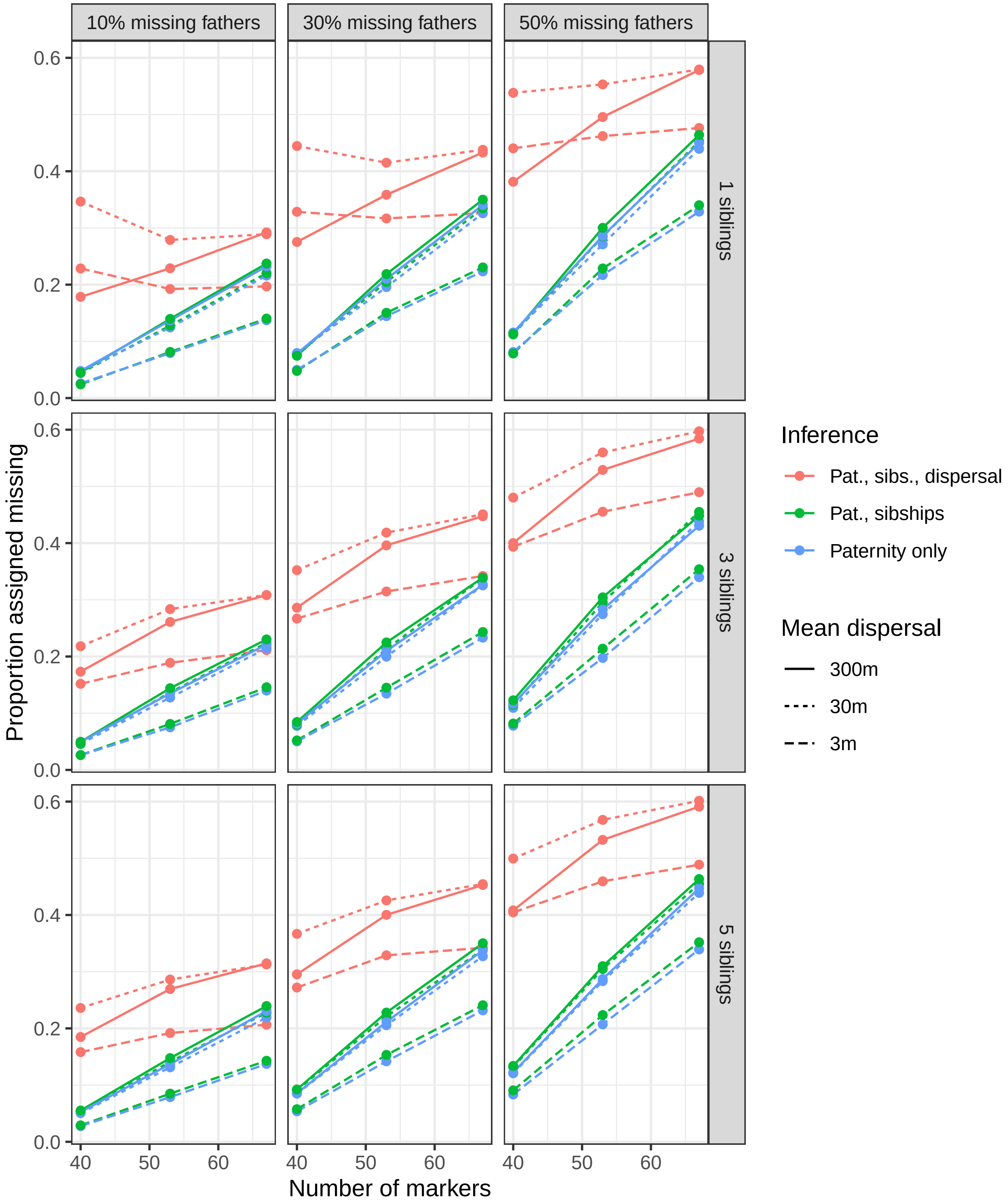


Figure S9: Proportion of offspring assigned to missing fathers in simulated data. Panels show the proportion of true fathers missing from the dataset used to infer mating events (columns) and the true full sibship size (rows). Values are the mean of 100 replicate simulations, and are shown for inference using paternity (Pat) only, paternity and sibships (sibs), and paternity, sibships and dispersal information.
